# poreCov - an easy to use, fast, and robust workflow for SARS-CoV-2 genome reconstruction via nanopore sequencing

**DOI:** 10.1101/2021.05.07.443089

**Authors:** Christian Brandt, Sebastian Krautwurst, Riccardo Spott, Mara Lohde, Mateusz Jundzill, Mike Marquet, Martin Hölzer

## Abstract

In response to the SARS-CoV-2 pandemic, a highly increased sequencing effort has been established worldwide to track and trace ongoing viral evolution. Technologies such as nanopore sequencing via the ARTIC protocol are used to reliably generate genomes from raw sequencing data as a crucial base for molecular surveillance. However, for many labs that perform SARS-CoV-2 sequencing, bioinformatics is still a major bottleneck, especially if hundreds of samples need to be processed in a recurring fashion. Pipelines developed for short-read data cannot be applied to nanopore data. Therefore, specific long-read tools and parameter settings need to be orchestrated to enable accurate genotyping and robust reference-based genome reconstruction of SARS-CoV-2 genomes from nanopore data. Here we present poreCov, a highly parallel workflow written in Nextflow, using containers to wrap all the tools necessary for a routine SARS-CoV-2 sequencing lab into one program. The ease of installation, combined with concise summary reports that clearly highlight all relevant information, enables rapid and reliable analysis of hundreds of SARS-CoV-2 raw sequence data sets or genomes. poreCov is freely available on GitHub under the GNUv3 license: github.com/replikation/poreCov.

## Introduction

It has been over a year since the WHO declared a pandemic related to SARS-CoV-2 on March 11, 2020. To date, this pandemic has caused over 112 million registered cases and more than 2.5 million deaths (March 2021). Over 190 countries have contributed to sequencing over 1.1 million publicly available SARS-CoV-2 genomes during this time, resulting in one of the greatest molecular surveillance efforts to date (data from GISAID; April 2021) (Elbe and Buckland-Merrett, 2017; Shu and McCauley, 2017). Thanks to global sequencing efforts, it is possible to track emerging variants, trace the evolution of the virus (Hodcroft et al., 2020; Brandt et al., 2021; Tang et al., 2021), and investigate potential escape mechanisms against vaccines at an early stage (Thompson et al., 2021; Zhou et al., 2021). In addition, the large amount of genomic data generated enables the development of simpler and faster rapid tests for the detection of variants of concern and characteristic mutations, e.g., for routine PCR screening in hospitals (Durner et al., 2021).

Illumina is still the most dominant approach for SARS-CoV-2 sequencing and genome reconstruction (Tab. 1). However, nanopore sequencing via tiled amplicon-based genome reconstruction (Quick et al., 2017; Yan et al., 2021, 2) is also heavily used and of particular interest, as laboratories can enter sequencing with a comparably low initial investment. While many laboratories can implement nanopore sequencing techniques rapidly, bioinformatics know-how, on the other hand, is still very scarce in science (Hodcroft et al., 2021). The observed shortage of bioinformaticians can be explained by the necessary overlap in different areas of biology, bioinformatics, and computer science. For the latter, in particular, there is an enormous shortage of skilled workers, making the development of robust and widely applicable software tools necessary (Perkel, 2019). This bottleneck in bioinformatics expertise concerns accurate genotyping and subsequent genome reconstruction of SARS-CoV-2 with direct impact to accompanying downstream analyses such as raw data and genome quality controlling, virus lineage determination, vaccine target surveillance, phylogenomics, and dynamic evolutionary observations and the various metrics behind them (Brandt et al., 2021; Hodcroft et al., 2021; Hufsky et al., 2021). Bioinformatics tools or workflows for similar tasks (e.g., reference-based genome reconstruction) are constantly being reinvented and developed independently by many laboratories. A broad application combined with user-friendly handling is rarely in focus. One reason for that is the additional development time and necessary maintenance. However, the usage of workflow management systems and containerization also allow smaller labs to provide reproducibility and, due to modularization, easy to maintain software tools (Perkel, 2019). A few workflows targeting the analysis of SARS-CoV-2 amplicon-based nanopore sequencing data already exist (e.g., github.com/nf-core/viralrecon, github.com/connor-lab/ncov2019-artic-nf), which are primarily aimed at a bioinformatics audience, not intended for public use or are over-complicated for biologists to use due to high level of necessary parameterization. While such features enable important workflow flexibility to support different sequencing protocols, they can be challenging to apply for a non-expert audience. However, due to the high sequencing workload during the pandemic, many laboratories urgently require bioinformatics support. We developed poreCov to diminish the bioinformatic entry barriers as much as possible and to allow reliable genotyping and reference-based genome reconstruction of SARS-CoV-2 samples in a high-throughput manner. While poreCov is optimized to analyze amplicon-based nanopore sequencing data following the widely used ARTIC protocol, the pipeline can also be applied to other primer sets and protocols. Reference-based genome reconstruction is the central focus of our pipeline, complemented by a group of downstream analyses for virus lineage determination and a comprehensive summary of results.

**Tab 1:** Sequencing technology used for SARS-CoV-2 surveillance. Data from GISAID screening 1,185,291 entries (date: April 2021). Entries for which multiple technologies were specified were counted once for each mentioned technology (increasing the total count by 1,085). 9,747 entries could not be assigned or were marked as unknown.

| **Sequencing technology** | **Number [n]** | **Proportion [%]** |
| --- | --- | --- |
| Illumina sequencing (Illumina) | 913,228 | 77.61 |
| Nanopore sequencing (Oxford Nanopore Technologies) | 202,503 | 17.21 |
| SMRT sequencing (Pacific Biosciences) | 30,447 | 2.59 |
| Ion Torrent (ThermoFisher) | 12,895 | 1.10 |
| DNBSEQ (MGI) | 12,658 | 1.08 |
| Sanger sequencing | 4,898 | 0.42 |

## Material and Methods

### General design

One of the main aspects of poreCov is to keep the bioinformatics part as simplistic as possible while still comprehensive and robust. Our development focused on easy installation, configuration, and execution on various computer systems with clear results and a minimal set of necessary input parameters. We developed customized scripts to collect and visualize all relevant results in a comprehensive summary report that comes as an intuitive HTML report as well as an Excel sheet (XLSX) and a tab-separated table (TSV) for direct programmatic access. To start an analysis with poreCov, only the bare necessity of inputs is required (where is the data located, what kind of input data is it, which primers are used). Other parameters are set to recommended default values but can still be changed by experienced users if necessary.

We use Nextflow (Di Tommaso et al., 2017) as a workflow manager giving poreCov the capabilities to be executed on most computational infrastructures from a laptop up to a compute cluster and cloud computing solutions. The native container support (Docker, Singularity) allows for comfortable, stable, and install-free execution of the various included tools without software dependency conflicts. Our containers are prebuilt, stored on docker hub (hub.docker.com), and automatically downloaded by poreCov if not already available on the system. This leaves Nextflow and either Docker or Singularity as the only dependencies needed to be installed for poreCov. If these dependencies are available, an initial installation and further updates of the workflow and the included tools can be performed with a single short command (‘nextflow pull replikation/poreCov’). When poreCov runs on a high-performance computing (HPC) cluster or cloud services, all resources (CPUs, RAM) for all processes and associated containers are pre-configured but can be adjusted via a user-specific configuration file when needed.

We use complete version control for poreCov from the workflow itself to every tool to achieve reproducible results. For this purpose, each of the more than 15 containers used is version-controlled based on their associated tool. Besides, each poreCov version can be accessed and executed individually, and the tool versions used during genome reconstruction and analysis are listed in the final report.

### Inputs

poreCov automatically adjusts the necessary workflow steps based on the provided input. As a result, a more flexible workflow execution is possible depending on the available data (Figure 1). As each sequencing lab has its own way for processing nanopore sequencing data, we allow for three different sequence inputs to start the genome reconstruction and variant calling steps: 1) a directory containing fast5 raw files (not basecalled), 2) an already basecalled directory containing fastq sequence files (e.g., the output from a MinIT or GridION). Additionally, 3) combined, single fastq file (one fastq file per sample) can be provided to simplify further the reanalysis of read data sets (e.g., from public repositories like ENA). Moreover, the genome reconstruction step is automatically skipped if already reconstructed genomes (fasta files) are provided as a direct input. By that, poreCov also allows to quickly and conveniently re-analyze previous genomes, e.g., to check for new clade or lineage assignments or to simply generate quality metrics and a summary report for a set of SARS-CoV-2 genomes. Optionally, a sample sheet can be provided to rename barcodes automatically and to simplify data analysis and file management post-analysis.

**Fig. 1:**
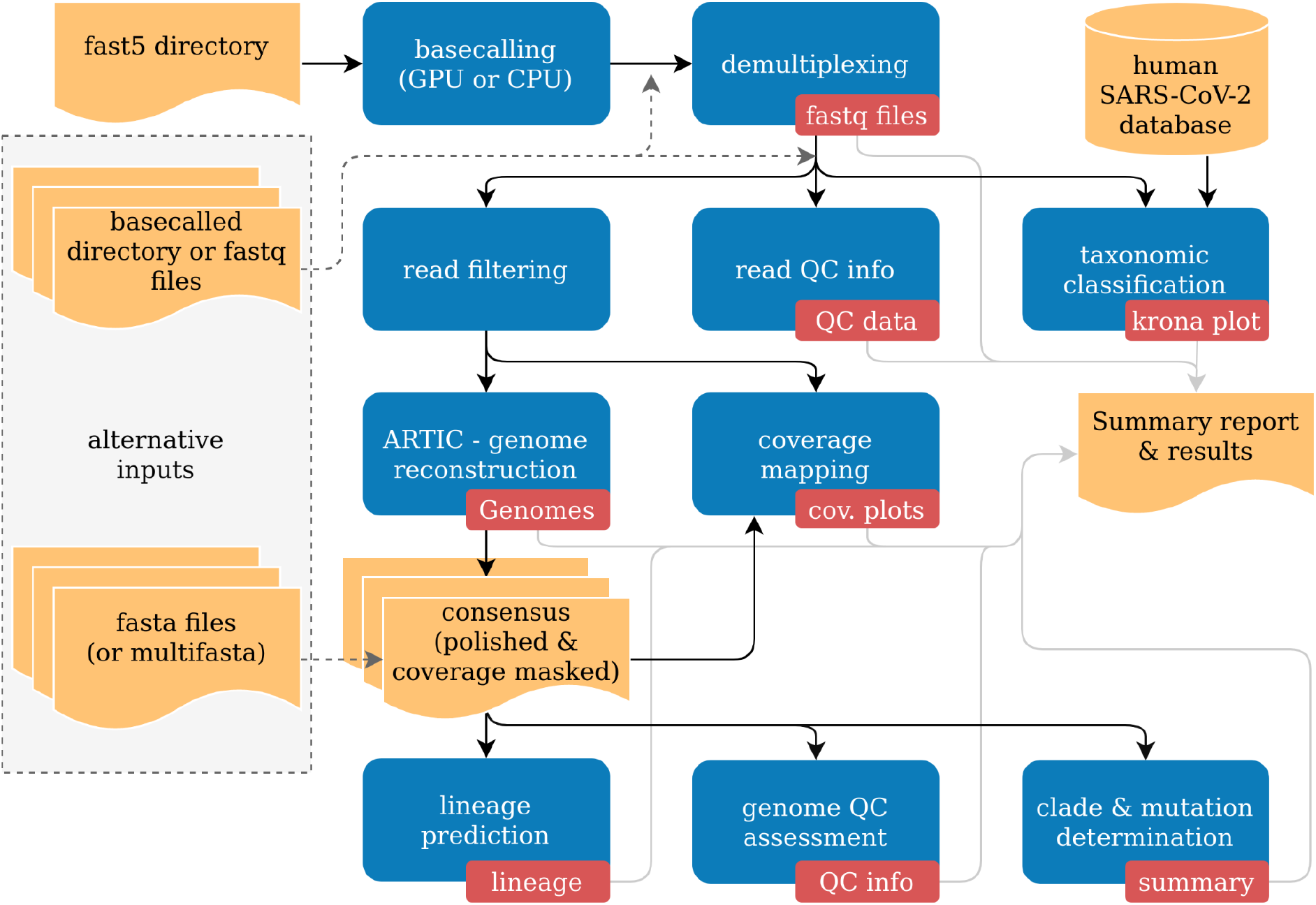
Simplified overview of the poreCov workflow. Results and inputs are colored in yellow. Workflow processes are colored blue, and the resulting information of a process is red.

### Included Analysis steps and tools

We heavily build poreCov around the widely used ARTIC protocol and pipeline (Quick et al., 2017). For the nanopore raw data (fast5) as an input, poreCov provides some necessary flexibility as the GPU basecaller Guppy (Oxford Nanopore Technologies) and the dependencies can be difficult to install. By default, poreCov assumes that either Docker or Singularity with GPU support can be used. However, users can switch to a local installation of Guppy or CPU-based basecalling or skip basecalling automatically by providing already basecalled reads (fastq).

Briefly, reads are prefiltered based on their length (default: 400-700 bp for ARTIC V3) followed by genome reconstruction via the ARTIC pipeline (github.com/artic-network/fieldbioinformatics). The ARTIC pipeline comprises tools and custom scripts for working with viral nanopore sequencing data generated from tiling amplicon schemes. General mappings are performed with minimap2 (Li, 2018), and intermediate file transformations are performed with SAMtools and BCFtools (Danecek et al., 2021). By default, genotyping is performed via medaka (github.com/nanoporetech/medaka) and Longshot (Edge and Bansal, 2019) but can also be done via nanopolish (github.com/jts/nanopolish) if fast5 data is provided. Regions with a low read coverage of 20X or less are masked by default. Furthermore, all reads are taxonomically classified via Kraken2 (Wood et al., 2019) against a pre-computed database comprised of the human GRCH38.p13 genome and all GISAID SARS-CoV-2 genomes as of February 2021 (available at doi.org/10.5281/zenodo.4534746) to unveil possible contamination or isolation issues from the wet lab. Krona plots (Ondov et al., 2011) are used to visualize the results. Reads are quality-controlled (read length distribution, quality scores, etc.) either via NanoPlot (De Coster et al., 2018) or pycoQC (Leger and Leonardi, 2019) depending on the user’s input. Additionally, all reads for each sample are mapped back against the Wuhan reference genome (Accession: NC_045512.2) using BWA-MEM (Li, 2013) to provide visual feedback of amplicon drop-outs, which are a common issue during the multiplex PCR step of the tiled amplicon sequencing protocol.

Each reconstructed and polished genome is further analyzed via pangolin (lineage determination; github.com/cov-lineages/pangolin), nextstrain (nextstrain clades, mutations, and deletions) (Hadfield et al., 2018), and president (calculating various genome quality metrics; gitlab.com/RKIBioinformaticsPipelines/president). Finally, all results are summarized in an HTML, TSV, and XLSX report to allow fast detection of low-performing samples, evaluation of negative controls, and an overview of virus lineages and detected mutations.

### Runtime & Resources

The execution of poreCov does not require a powerful computer, and only four threads (two CPUs) are sufficient for this purpose. However, the analysis will take longer (Tab. 2). poreCov shows its strengths on larger workstations or HPCs with full parallelization through the used workflow management system. By default, poreCov utilizes all available threads and tries to run at least four jobs in parallel for the analysis. If necessary, this behavior can be individually controlled and fine-tuned via the cores option flag (Tab. 2).

**Tab. 2:**
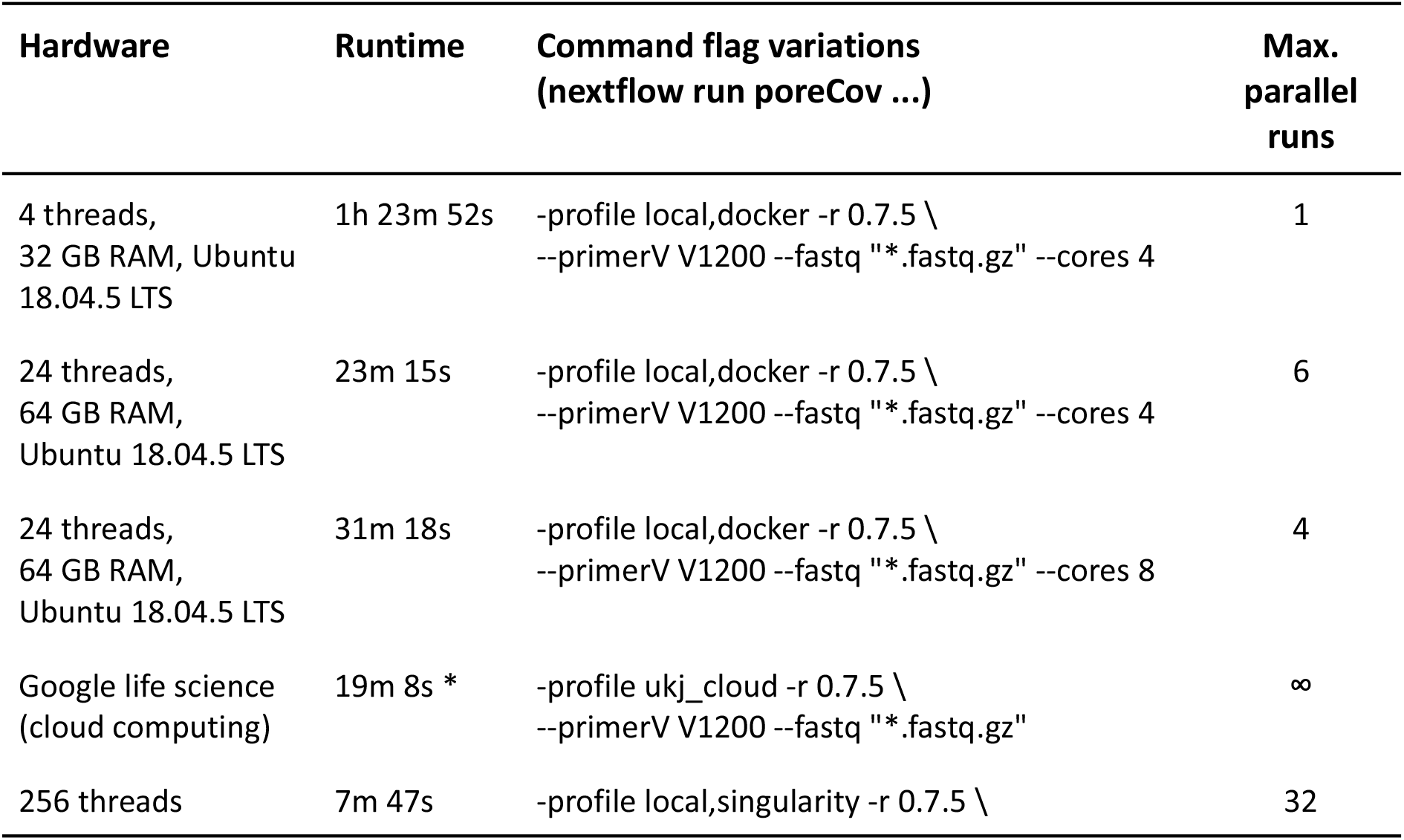
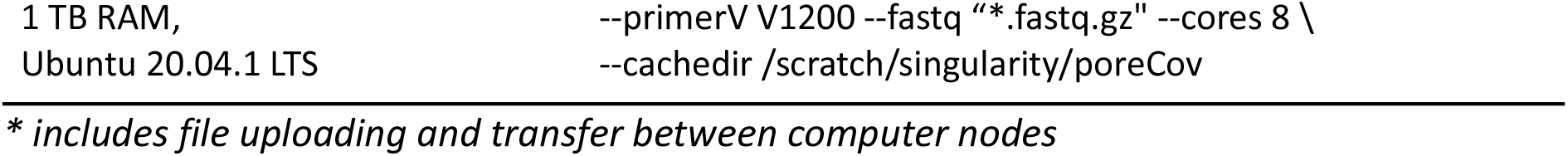
poreCov time to results (runtime) for different hardware configurations using ten read files as input (fastq files). The database for Kraken2 and images for Docker/Singularity were pre-downloaded and already available on the different systems. The test runs were performed with sequencing data from the ARTIC 1200 bp amplicon protocol.

## Results & Discussion

### Result reporting

Quick and easy-to-understand result interpretation is a crucial challenge for labs that want to analyze hundreds of SARS-CoV-2 genomes or more in short periods. To achieve this, poreCov summarizes all the necessary results besides the reconstructed genomes in a final report file (see Figure 2).

**Fig. 2:**
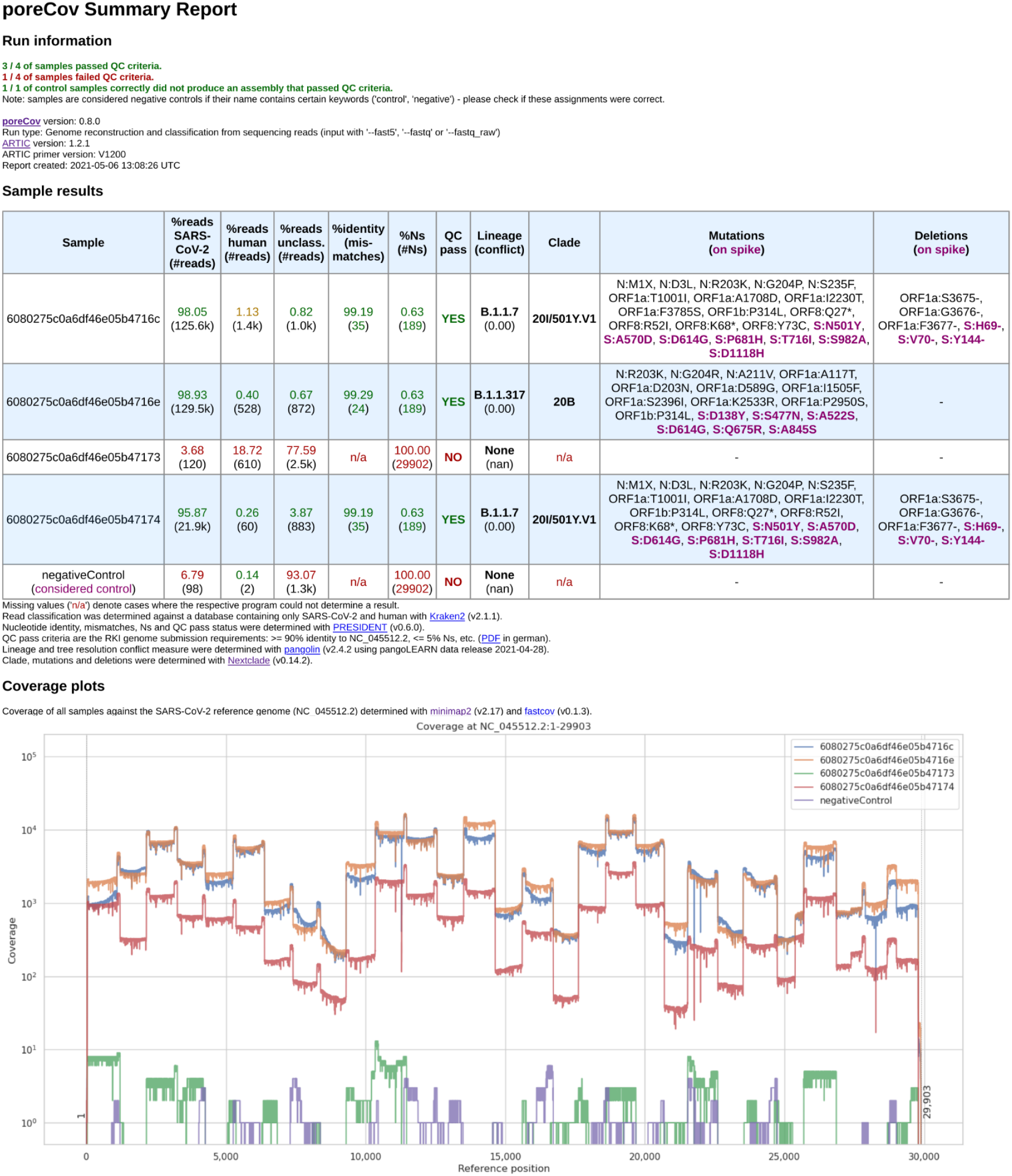
Final report generated by poreCov after each run. Four samples and one negative control were analyzed using 1200 bp amplicon primers. The first part of the report, the run information, summarizes the overall performance of the sequencing run in a short form across all samples. The second section, the sample results, summarizes all relevant information for each sample in tabular form. The coverage plots are presented in batches of 6 samples, and additional charts are automatically added for larger runs.

The first part of the report is a table containing the relevant information regarding contaminations (amount of human, viral and unclassified reads). The second part includes the overall quality of the genomes (percent identity to the Wuhan genome, amount of ambiguous nucleotides). All these features are summarized into “QC pass” if genomes have >= 90% identity to the Wuhan strain (NC_045512.2) and <= 5% ambiguous nucleotides (Ns). The last columns of the table highlight the determined lineages based on the pangolin and nextclade nomenclatures and include information about substitutions and deletions based on the nextclade alignments. Any substitutions or deletions in the spike protein are highlighted. Furthermore, clear color codings (green, yellow, red) are used to quickly allow the user to visually screen for issues or problems that might have arisen throughout the genome reconstruction and analysis steps. The last part of the report is coverage plots shown in batches of six samples to rapidly assess why a certain sample did not yield a genome that meets the QC criteria or in which regions the coverage was low (e.g. amplicon drop-outs due to primer dimerization (Itokawa et al., 2020)).

So far, poreCov was used and evaluated on more than 600 SARS-CoV-2-PCR-positive samples analyzed at the Jena University Hospital, Thuringia (Germany) and over 400 samples at the University Hospital Regensburg, Bavaria (Germany). In addition, the pipeline is routinely used for genome reconstruction at Germany’s Public Health Institute and analyzed between February 23rd and May 5th, 2021, 1,012 samples sequenced on MinION or GridION machines.

### SARS-CoV-2 genomic surveillance via nanopore sequencing: challenges and pitfalls

During our work on the poreCov pipeline, we identified several common challenges and pitfalls that arise, particularly during library preparation and sequencing, which can significantly impact downstream bioinformatics analyses. Here, we share such common issues in the context of SARS-CoV-2 nanopore sequencing and how poreCov can help to identify and subsequently avoid these.

#### Sample-Barcode bleeding

As nanopore sequencing is a molecule-based approach, it is easy to “over- or underload” cDNA molecules for library preparation. This overloading often happens if too many “super small” fragments (< 100 nucleotides) are present in the sample. Molecule-overloading usually leads to unspecific barcode ligation during the adapter ligation step after barcode pooling. Free barcodes from samples with a low amount of DNA molecules might ligate to non-barcoded DNA from samples with a highly overloaded number of molecules. A negative control can visualize this issue (poreCov highlights negative controls in the report if specified). Furthermore, using only reads with barcodes present on both ends mitigates the problem (the default setting for poreCov). If sample bleeding occurred, a false-positive negative control usually inherits a similar SNP proportion pattern as the barcoded samples used on the same run (see Fig. 3). This can be visually inspected by opening the .bam file in the result dir “2.Genomes” via, e.g., Unipro UGENE (Okonechnikov et al., 2012) or IGV viewer (Robinson et al., 2011).

**Fig. 3:**
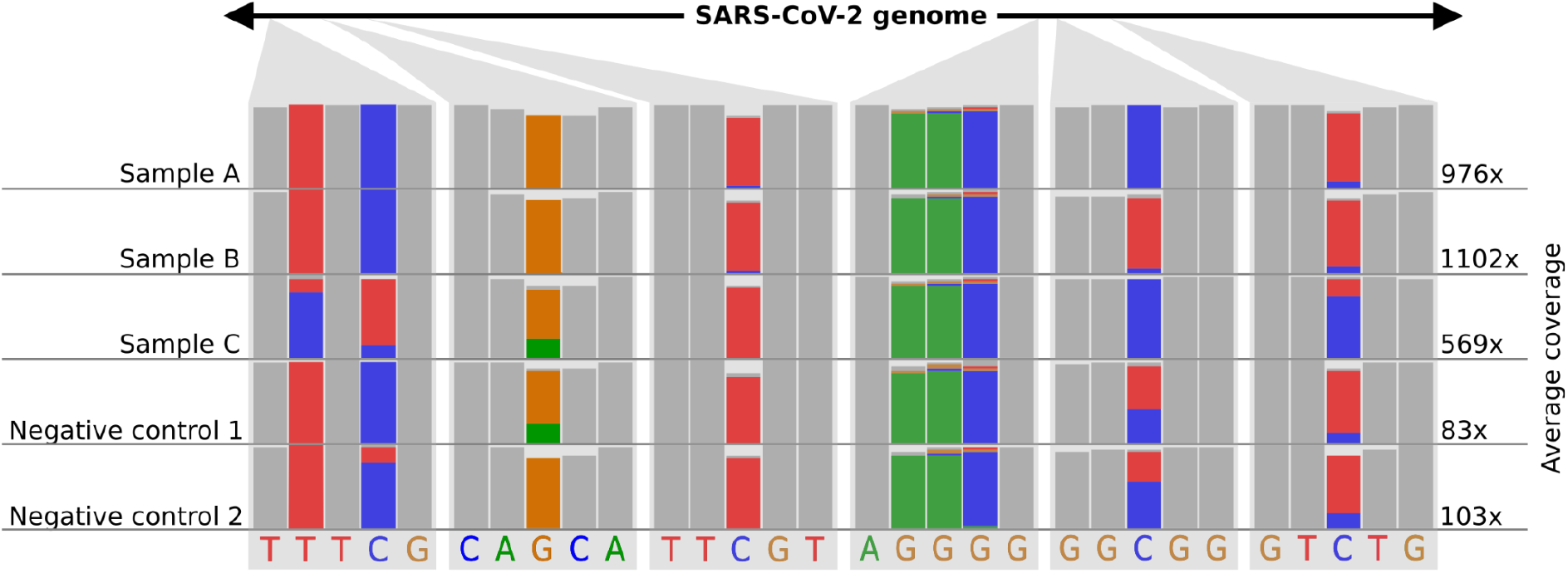
Example of how sample-barcode bleeding can be identified by inspecting SNP proportions of false-positive negative controls in mapping (.bam) files. Library preparation for samples A to C was performed with too much DNA. Negative controls follow the same library preparation steps as the samples A-C but with pure water instead of viral amplicon DNA. Additionally, the library preparation for the negative controls was performed in a separate lab and with new reagents up until the barcode pooling. Negative controls indicate SNP proportions that roughly reflect sample A to C considering their average coverage. The schematic figure is adapted from IGV.

#### Missing amplicons

A common problem after SARS-CoV-2 genomes reconstruction are long stretches of ambiguous nucleotides due to amplicon dropouts because of the high number of primers during the multiplex PCR. An alternative to the commonly used ARTIC V3 amplicon protocol uses longer 1200 bp amplicons to decrease the overall number of primers used. Longer amplicons usually lead to higher overall throughput (total bases) due to the longer read median. poreCov supports both primer sets (the standard 400 bp amplicons and longer 1200 bp amplicons) and visualizes each sample’s coverage plot to identify errors during the multiplex PCR step (see Fig. 2). A possible solution is to increase primer concentration for these specific “low coverage” regions. Additionally, it is possible to just re-amplify the missing regions via a cheap flongle flowcell, combine both read sets and re-construct and re-analyze via poreCov. For this, poreCov screens each genome and reports back which minimal set of primers would be needed to amplify only regions with ambiguous bases.

#### Many ambiguous bases

Besides missing amplicons (see above), the genome may contain many masked regions even though DNA concentrations and the number of sequenced reads were high. Most labs use direct patient material from swabs to sequence SARS-CoV-2 genomes to quickly return results without delay. Therefore, the sampling quality directly influences the sequencing run as any given sample could still have more human RNA/DNA than viral after isolation or if some of the isolation steps went wrong. This issue can be easily identified via poreCov as the amount of viral and human reads for each sample are summarized. Typical samples contain 90 % or more viral reads after sequencing, and decreasing proportions might fail the genome reconstruction step or result in low-quality genomes (see Fig. 2, columns: “%reads”).

#### Many unclassified barcodes

When using only reads with barcodes on both ends, this might lead to many reads without a barcode determination if something went wrong during the barcoding step in the wet lab. The number of unclassified barcodes can be visually inspected via the pycoQC report in the “1.Read_quality” dir. A possible solution is to deactivate both end requirements via poreCov’s ‘--one_end’ option flag. However, this can lead to the “strain bleeding” problem described above and is also not recommended (artic.network/quick-guide-to-tiling-amplicon-sequencing-bioinformatics.html). Therefore, it is strongly recommended to adjust the wet lab procedures as longer ligation times and lower DNA mol concentrations usually resolve this.

#### Staying up to date with lineage classifications

Lineage classification via pangolin is widely used and continuously updated and corrected based on the current pandemics’ prevalent virus variants. Due to this, the pangolin lineage assignment step in poreCov needs to be updated regularly. We ensure this via automatic, version-controlled, and tested container creation. Valid containers are automatically pushed and stored on hub.docker.com/r/nanozoo/pangolin. poreCov can check what is the latest available version for lineage classification before each run. If up-to-date lineage assignments are crucial, this feature can be manually activated and is deactivated by default to ensure the stability of the pipeline and stable offline usage. The pangolin version and the pangoLEARN version (the trained model for classification) are both noted on the final report to ensure reproducibility.

## Conclusion

Nanopore sequencing represents an important pillar in the sequencing of SARS-CoV-2 genomes and thus in the molecular surveillance of novel viral variants and outbreak events of the global pandemic. However, due to its novelty and rapid development, nanopore sequencing also poses downstream challenges and pitfalls for accurate bioinformatic analysis and reproducibility. Many laboratories have ramped up nanopore sequencing, but are in dire need of bioinformatics support given the high volume of global sequencing during the SARS-CoV-2 pandemic, especially when hundreds of samples need to be processed recurrently. Here we present poreCov, a high-throughput software pipeline for reliable genotyping and reference-based genome reconstruction of SARS-CoV-2 samples based on nanopore tiled amplicon sequencing. The pipeline is built around the widely used ARTIC protocol and integrates a comprehensive summary report with a clear result structure, allowing a quick assessment of the genome reconstruction performance. Clear color highlighting allows straightforward result interpretation even for large sequencing batches. Typical pitfalls during routine sequencing can be, therefore, quickly spotted and solved. In addition, the user can access all of the intermediate data via a well-structured result directory for further analysis or troubleshooting. We ensure a high degree of reproducibility and stability by utilizing Nextflow and virtualizing each tool through containers. We made sure that poreCov can run with the minimal required inputs and robust default parameters to diminish the bioinformatic entry barriers and enhance SARS-CoV-2 genome reconstruction and analysis during the pandemic.

## Availability and future directions

The poreCov workflow is fully modularized to allow straightforward integration of additional processes, update tools, and add, if necessary, new primer schemes. As the primary framework utilizes the ARTIC protocol for genome reconstruction (as before also applied for, e.g., ZIKA and EBOV) (Quick et al., 2017), it can be easily adjusted to work for other viruses and potential further outbreaks. By that, “sub-workflows” of the pipeline can be automatically deactivated or conditionally activated based on the input data to avoid convoluting the workflow with parameters that need to be set by the user. Because poreCov is productively used at the public health institute in Germany, the SARS-CoV-2 side of the workflow will be continuously updated and extended if the necessity arises.

## Acknowledgment

We thank Stephan Fuchs and Oliver Drechsel for creating and providing the Kraken2 database, used in gitlab.com/RKIBioinformaticsPipelines/ncov_minipipe, and Kathrin Trappe for feedback on running and testing the pipeline continuously.

We thank all the people on GitHub that wrote poreCov issues and helped to identify and resolve bugs.

We thank the GISAID team for providing metadata about the sequencing technologies used for SARS-CoV-2 sequencing.

